# Microcirculatory dysfunction associates with neurovascular uncoupling in peri-ischemic brain regions after ischemic stroke

**DOI:** 10.1101/2022.08.26.505245

**Authors:** Christian Staehr, John T. Giblin, Eugenio Gutiérrez-Jiménez, Halvor Guldbrandsen, Jianbo Tang, Shaun L. Sandow, David A. Boas, Vladimir V. Matchkov

**Affiliations:** Department of Biomedicine, Aarhus University, Aarhus, Denmark; Neurophotonics Center, Department of Biomedical Engineering, Boston University, Boston, MA, USA; Center of Functionally Integrative Neuroscience, Institute for Clinical Medicine, Aarhus University, Aarhus, Denmark; Department of Biomedical Engineering, Southern University of Science and Technology, Shenzhen, China; Discipline of Biomedical Science, School of Health and Sports Science, University of the Sunshine Coast, Maroochydore, QLD, Australia; Faculty of Medicine, Centre for Clinical Research, The University of Queensland, Brisbane, QLD, Australia

## Abstract

**Background:** Despite recanalization after ischemic stroke, neurovascular coupling, i.e., the local hyperaemic response to neuronal activity, is impaired in peri-ischemic brain regions. Reduced neurovascular coupling may contribute to neurological deterioration over time. The mechanism underlying dysfunctional neurovascular coupling following stroke is largely unknown.

**Methods:** Mice implanted with chronic cranial windows were trained for awake head-fixation prior to experiments. One hour occlusion of the anterior middle cerebral artery branch was induced using single vessel photothrombosis. Cerebral perfusion and neurovascular coupling were assessed by optical coherence tomography and laser speckle contrast imaging. Capillaries and pericytes were studied in perfusion-fixed tissue by labelling lectin and platelet-derived growth factor receptor β.

**Results:** Arterial occlusion induced on average 11 spreading depressions over one hour associated with substantially reduced blood flow in the peri-ischemic cortex. Approximately half of the capillaries in the peri-ischemic area were no longer perfused 3 and 24 hours after reperfusion, which was associated with constriction of an equivalent proportion of peri-ischemic capillary pericytes. The capillaries in the peri-ischemic cortex that remained perfused showed increased prevalence of dynamic flow stalling. Whisker stimulation led to reduced neurovascular coupling responses in the sensory cortex corresponding to the peri-ischemic region 3 and 24 hours after reperfusion.

**Conclusion:** Arterial occlusion led to constriction of pericytes in the peri-ischemic cortex associated with long-lasting microcirculatory failure. This reduced capillary capacity may, at least in part, underlie impaired neurovascular coupling in peri-ischemic brain regions after stroke and reperfusion.

## Introduction

In the central nervous system, there is an accurately balanced coupling between neuronal tissue activity and function of the supplying vasculature. Neurovascular coupling ensures rapidly increased supply of oxygen and nutrition to active brain regions (Nippert et al., 2018). The communication is believed to be transmitted through release of vasoactive substances from the active neuronal tissue, which in turns dilate parenchymal arterioles and thus increases blood flow (Longden et al., 2017; Staehr et al., 2020). Furthermore, the capillary circulation plays an important role in the modulation of neurovascular coupling. It has been suggested that pericytes associated with cerebral capillaries contain smooth muscle actin and are contractile (Bandopadhyay et al., 2001; Erdener et al., 2022; Hall et al., 2014), with the related mechanism preferentially shunting capillary blood flow for specific tissue perfusion. Nevertheless, whether pericytes actively regulate capillary diameter in response to neuronal activity and thus contribute to neurovascular coupling remains controversial (Fernández-Klett et al., 2010; Hall et al., 2014; Hill et al., 2015; Kisler et al., 2017).

Impaired neurovascular coupling, which may contribute to neurological deterioration over time (Kisler et al., 2017), has been reported in patients following ischemic stroke (Krainik et al., 2005; Lin et al., 2011; Pineiro et al., 2002). Importantly, these studies reported that reduced neurovascular coupling is also observed in brain regions apart from the ischemic core. Hence, it has been speculated that impaired neurovascular responses in stroke patients were caused by a pre-existing diffuse vascular pathology (D’Esposito et al., 2003; Pineiro et al., 2002). The present study aimed to address this unanswered question by comparing neurovascular responses in the peri-ischemic area before cerebral ischemia and after ischemia-reperfusion. We hypothesized that impaired neurovascular coupling in the peri-ischemic area is caused by disrupted microcirculation. We tested the hypothesis in awake mice by performing a thorough characterization of cerebrovascular function in the peri-ischemic area before, during, and after cerebral ischemia. By studying the dynamic changes in cerebral blood flow during arterial occlusion and after ischemia-reperfusion, a unique mechanistic insight into the cerebrovascular events in ischemic stroke was obtained. With state-of-the-art imaging techniques, a mechanism that may underlie wide-spread neurovascular uncoupling in stroke patients was uncovered.

## Methods

All experiments were performed in awake mice unless otherwise stated, with all data included in the analysis.

### Experimental animals

All experiments were approved by the Institutional Animal Care and Use Committees at Boston University. Experiments were conducted following the Care and Use of Laboratory Animals guidelines and reported in accordance with the ARRIVE (Animal Research: Reporting in vivo Experiments) guidelines. Female C57BL/6 mice (Jackson Laboratory, Bar Harbor, ME, USA) were used. Mice were 18 weeks old when chronic cranial windows were prepared and between 24 and 26 weeks old when the stroke-reperfusion protocol commenced. Mice were housed under a 12:12 light/dark cycle, and food and water were provided ad libitum.

### Surgical procedure and habituation training

Four hours prior to the surgical procedure, 4.8 mg/kg dexamethasone (4 mg/mL) was administered i.p. to minimize cerebral tissue swelling during surgery. Two hours prior anaesthesia, 0.5 mg/kg slow-release buprenorphine (1 mg/mL) and 5 mg/kg meloxicam (5 mg/mL in 0.9% NaCl) was injected subcutaneously. Body temperature was maintained at 37°C during surgery and controlled with a rectal probe. Surgery was performed under isoflurane anaesthesia (2–3% induction, 1–2% maintenance, in 1 L⁄ min oxygen). Craniotomy over one hemisphere was performed. Dura mater was left intact. The brain surface was covered with a piece of curved glass (LabMaker, Berlin, Germany) that was shaped to fit the cranial window and sealed with dental acrylic and instant super adhesive glue. A circular aluminium bar was attached to the skull around the cranial window for head fixation during experiments. Meloxicam (5 mg/kg) was administered every 24h three days following surgery. Mice were allowed to recover from surgery for at least 14 days before starting the habituation training. Mice were gradually habituated to longer periods of head-fixation over seven training sessions starting at a duration of 10 min and ending at 1.5 h. While head-fixed, mice were able to readjust body position. After each habituation session, mice were rewarded with sweetened milk.

### Photothrombosis

Single vessel photothrombosis was performed as described previously (Sunil et al., 2020) to occlude the anterior middle cerebral artery (MCA) branch while causing minimal damage to the surrounding parenchyma (Fig. 1A-B). Under brief isoflurane sedation (1% in 1 L⁄ min oxygen), 100 μl of Rose Bengal (15 mg⁄ml in 0.9% NaCl; Sigma Aldrich, St. Louis, MO, USA) was injected retro-orbitally. Rose Bengal is activated with green light. Therefore, blue light was used to visualize the vasculature with a multispectral camera and thus did not lead to photoactivation. The green laser diode (520 nm) was tuned to a post-objective power of 0.6 mW and focused to a diameter of 6 μm. Real-time laser speckle contrast imaging (LSCI) was used to detect when the target vessel was occluded. When no blood flow was observed, the green laser was left on for another 2 min and then turned off. In the case of spontaneous reperfusion during the 1 h of occlusion, the green laser was turned on until the vessel was occluded again. Collateral blood flow supply can prevent single vessel occlusion from having a significant reduction in blood flow in mice and thus resulting in the formation of an ischemic core (Sigler et al., 2008; Sunil et al., 2020). Therefore, a collateral artery was targeted if it appeared after occlusion and supplied the perfusion field for the occluded MCA branch. Collateral blood supply was targeted in half of the mice.

**Figure 1.**
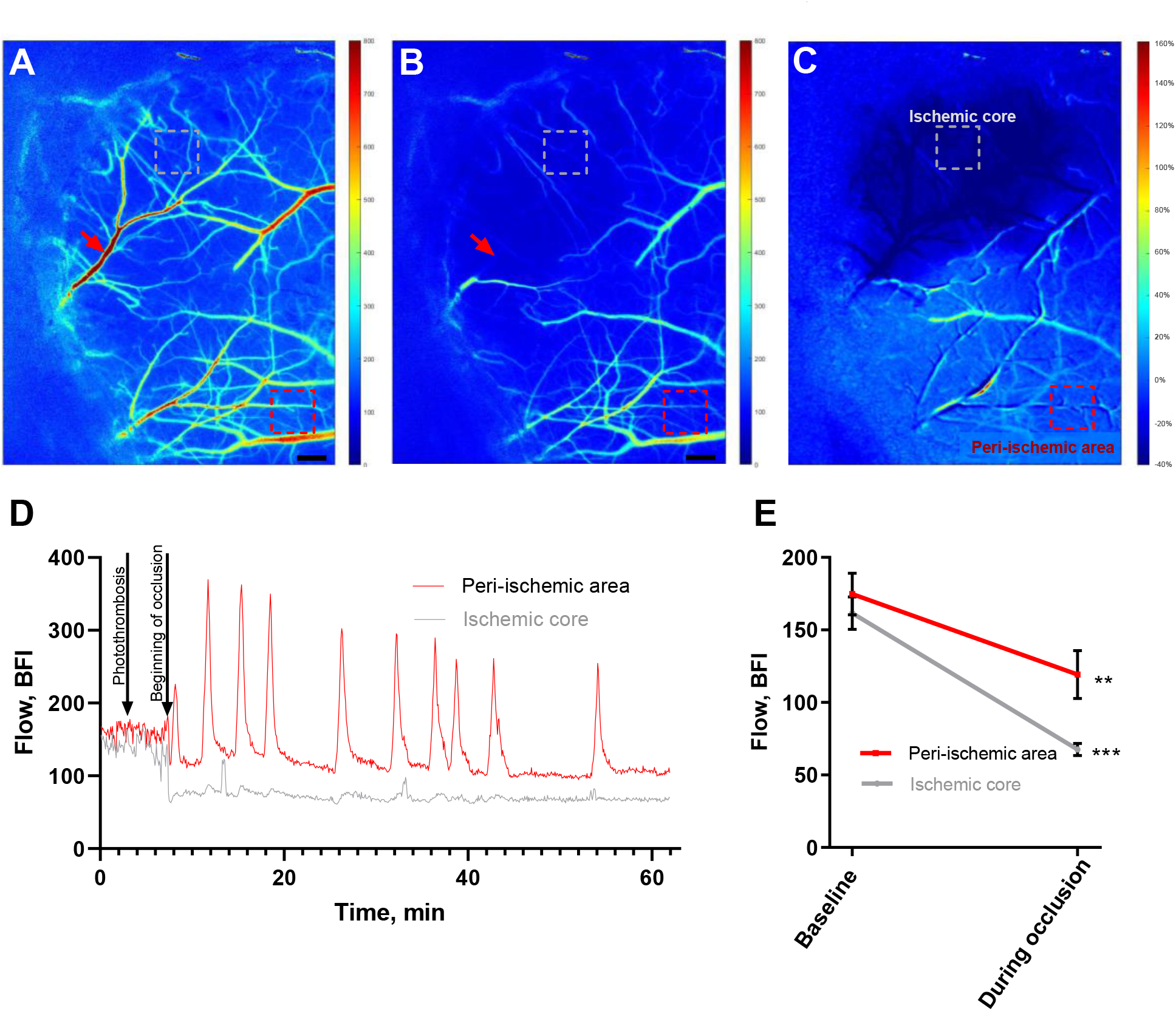
Cerebral ischemia induced by single vessel photothrombosis of the middle cerebral artery. Representative laser speckle contrast images (LSCI) of the affected hemisphere in the awake mouse at baseline (A) and during occlusion of the anterior middle cerebral artery (MCA) branch (B). Bars in (A, B) correspond to 300 μm. Image of the ratio of LSCI before vs. during the occlusion is shown in (C). Red arrows in (A, B) show the focus point for the green laser beam (Ø = 6 μm) used for photothrombosis of the anterior MCA branch. Dotted boxes in (A-C) indicate the region of interest in the ischemic core (grey) and in the peri-ischemic area (red). Representative traces from these two regions show the changes in blood flow (BFI: blood flow index) during the MCA occlusion (D). MCA occlusion was associated with drop in blood flow in the ischemic core and, to a lesser extent, in the peri-ischemic area (E). Blood flow during occlusion was compared to baseline using two-tailed paired t-test. **, *** indicate *P* < 0.01, 0.001; *n* = 6.

### Optical coherence tomography

A spectral-domain optical coherence tomography (OCT; Telesto III, Thorlabs, Newton, NJ, USA) system with a 1310 nm center wavelength and a bandwidth of 170 nm was used for this study. The axial spatial resolution was 3.5 μm in the brain, which was determined by the light source. A 10× objective NA = 0.28 (Mitutoyo, Kawasaki, Japan) was used to image the cerebral microcirculation, resulting in a lateral spatial resolution of 3.5 μm in the brain (Erdener Ş et al., 2019). The focused beam was adjusted at a depth of around 150-200 μm, which enabled acquisition of blood flow signal down to ∼300 μm from the brain surface. The blood vessels were imaged with OCT angiography (OCTA) (Srinivasan et al., 2010), which detects the intrinsic dynamic contrast of the moving red blood cells by taking the difference of repeated OCT B-scans. A larger 3 × 3 mm field of view region and a smaller 600 × 600 μm field of view region were obtained at the baseline, 3h and 24h, respectively, after reperfusion. For the larger field of view, five angiograms were obtained and averaged. For the smaller field of view, 90 sequential OCT angiograms were obtained over a period of approximately 10 min. Maximum intensity projections over a range of 90 μm were extracted for analysis in the depth of 210-390 μm from the brain surface. This range was selected because of the dense capillary network at this depth.

Dynamic capillary flow stalling was defined as sudden intensity drop and thus disappearing of a capillary segment in ≥ 1 of the 90 consecutive smaller 600 × 600 μm field of view angiograms. Capillary segments that appeared at any of the 90 angiograms were included when counting perfused capillary segments. Previous analysis of dynamic flow stalling of capillaries with the use of the same approach showed that day-to-day variations between the measurements from the same region of interest were minimal (Erdener Ş et al., 2019). Counting of capillaries was done manually using ImageJ software (NIH, Bethesda, MD, USA).

Phase-resolved Doppler OCT (prD-OCT) was recorded with 25 repeated A-lines to assess the axial blood velocities in penetrating cortical blood vessels (Tang et al., 2017; Tang et al., 2019). PrD-OCT was done in the same region of interest as the smaller field-of-view angiogram. Capillaries with diameters ranging between 3-6 μm penetrating the cortex in a vertical direction were marked manually for each vessel in a single depth. Capillaries were selected over a depth range of 180-300 μm. To minimize background noise, a threshold was set to exclude velocities slower than 0.1 mm/sec. The OCTA and prD-OCT MATLAB codes are available at https://github.com/BUNPC/OCTA.

### Laser speckle contrast imaging and air-puff whisker stimulation

Coherent near-infrared light was delivered by a laser diode (785 nm, LP785-SAV50, Thorlabs Inc.). The diode was controlled by current and temperature controllers (LDC210C and TED200C, respectively; Thorlabs Inc.). Backscattered light was recorded using a 2x lens (TL2X-SAP, Thorlabs Inc.) mounted on a CMOS camera (acA2040-90um, Basler AG, Ahrensburg, Germany). A 650 nm long pass filter prevented the 520 nm photothrombosis laser from affecting the laser speckle contrast imaging (LSCI). LSCI data were analyzed by applying the commonly used model of arbitrary blood flow index calculated by 1/K^2^ where K is contrast (Boas and Dunn, 2010). Baseline and follow-up recordings were made over 180 sec with a framerate of 194 Hz and a resolution of 1024×512 pixels. During the 60 min occlusion, LSCI frame rate was set to 5 Hz and resolution to 1800×1200 pixels. The pixel size for all LSCI recordings was 3 μm. The LSCI data were processed using temporal speckle contrast analysis, where 25 frames were used to calculate contrast. A segmentation algorithm (Postnov et al., 2016) was used to analyse single vessel flow velocity as previously described (Staehr et al., 2019). Arbitrary flow velocity was based on the average laser speckle contrast value in the lumen of the vessel segment.

Whisker stimulation was done using air-puff. Five sec of whisker stimulation was preceded by 5-sec baseline and followed by 20-sec without stimulation. This 30-sec protocol was repeated 20 times, and the relative change in blood flow to each baseline period was assessed by LSCI. In whisker stimulation experiments, the frame rate was 40 Hz, and data were processed using spatial speckle contrast analysis to get a high temporal resolution. Custom-made MATLAB scripts used to process and analyse LSCI data are available at https://github.com/BUNPC/laserSpeckleImaging.

### Labelling of capillaries and pericytes

For structural analyses, mice were perfusion fixed through the left ventricle under 3% isoflurane with 4% formaldehyde in Dulbecco’s CaCl_2_ and MgCl_2_ free phosphate buffered saline (PBS). Brains were subsequently postfixed for 24h in 4% formaldehyde in PBS at 21°C and afterwards stored in PBS at 4°C. The fixed brains were embedded in paraffin and sectioned in 5 µm slices at the level of the peri-ischemic sensory cortex, approximately corresponding to the region of interest for OCTA assessment of capillary flow. Sections including both hemispheres were incubated for 16 h in primary antibody (1:500 rabbit anti-PDGFRβ, #ab32570, Abcam, Cambridge, UK) at 4°C followed by 2h incubation in secondary antibody (1:500 goat anti-rabbit IgG, Alexa Fluor 488, #ab150077, Abcam) at 21°C. Lipofuscin autofluorescence was reduced using TrueBlack Autofluorescence Quencher (1:20, Biotium, Fremont, CA, USA) for 60 sec. The whole sections were imaged using Olympus VS120 slide scanner (Olympus, Tokyo, Japan) at ×40 magnification. After imaging the pericytes, the samples were prepared for lectin labelling: Samples were treated for 5 min with pepsin (4 mg/mL PBS) at 37°C prior to 2h lectin incubation at 21°C (1:100 Lycopersicon Esculentum (Tomato) Lectin, DyLight™ 649, DL-1178-1 (10 µg/mL), Vector Labs, Newark, CA, USA). Nuclei were stained with 300 nM DAPI before mounting. The slide scanning was repeated, and the images with the two different labellings of the same tissue were registered and overlayed using Control Points Registration in Matlab (ver. R2022a, Natick, MA, USA). Capillary diameter was assessed automatically in ImageJ (National Institutes of Health) using VasoMetrics (McDowell et al., 2021).

### Data analyses

MATLAB R2022a and GraphPad Prism software (v.9.3.1) were used for graphing and statistical analyses. Data are summarized as the mean value ± SEM of the sample group. Significant differences between means were determined by either one-way ANOVA, mixed-effects analysis, or two-way ANOVA where appropriate, followed by correction for multiple comparison. A probability (*P*) level of <0.05 was considered significant.

## Results

### Arterial occlusion resulted in spreading depressions

With the use of LSCI it was ensured that single vessel photothrombosis led to 60 min no-flow in the anterior branch of MCA (Fig. 1A-B). This outcome was associated with 57 ± 3% drop in perfusion of the downstream cortex (Fig. 1C-E). Blood flow in the peri-ischemic area corresponding to the whisker sensory cortex was reduced by 33 ± 6% during occlusion (Fig. 1D). All six mice showed spontaneous reperfusion of the targeted artery within 3h. Within minutes after onset of the occlusion, spreading depressions propagated from the peri-ischemic area to the rest of the hemisphere (Suppl. Video 1 and Fig. 2A, B). Spreading depressions were observed during occlusion in all six mice. On average, 11 spreading depressions were observed during the 60 min occlusion period (Suppl. Table 1). The propagation velocity of the spreading depressions was 3-5 mm/min (Suppl. Table 1). For the first observed spreading depression, a period of hypoperfusion preceded a subsequent period of hyperperfusion followed by sustained hypoperfusion (Suppl. Video 1 and Fig. 2C). The duration of the initial hypoperfusion was shorter than the subsequent hyperperfusion (Suppl. Table 1). After the first spreading depression, blood flow was globally reduced. During this global hypoperfusion, several spreading depressions propagated and were associated with waves of hyperperfusion (Fig. 2D and Suppl. Table 1). The initiation of spreading depressions was not associated with the photothrombosis laser being on nor with spontaneous reperfusion.

**Figure 2.**
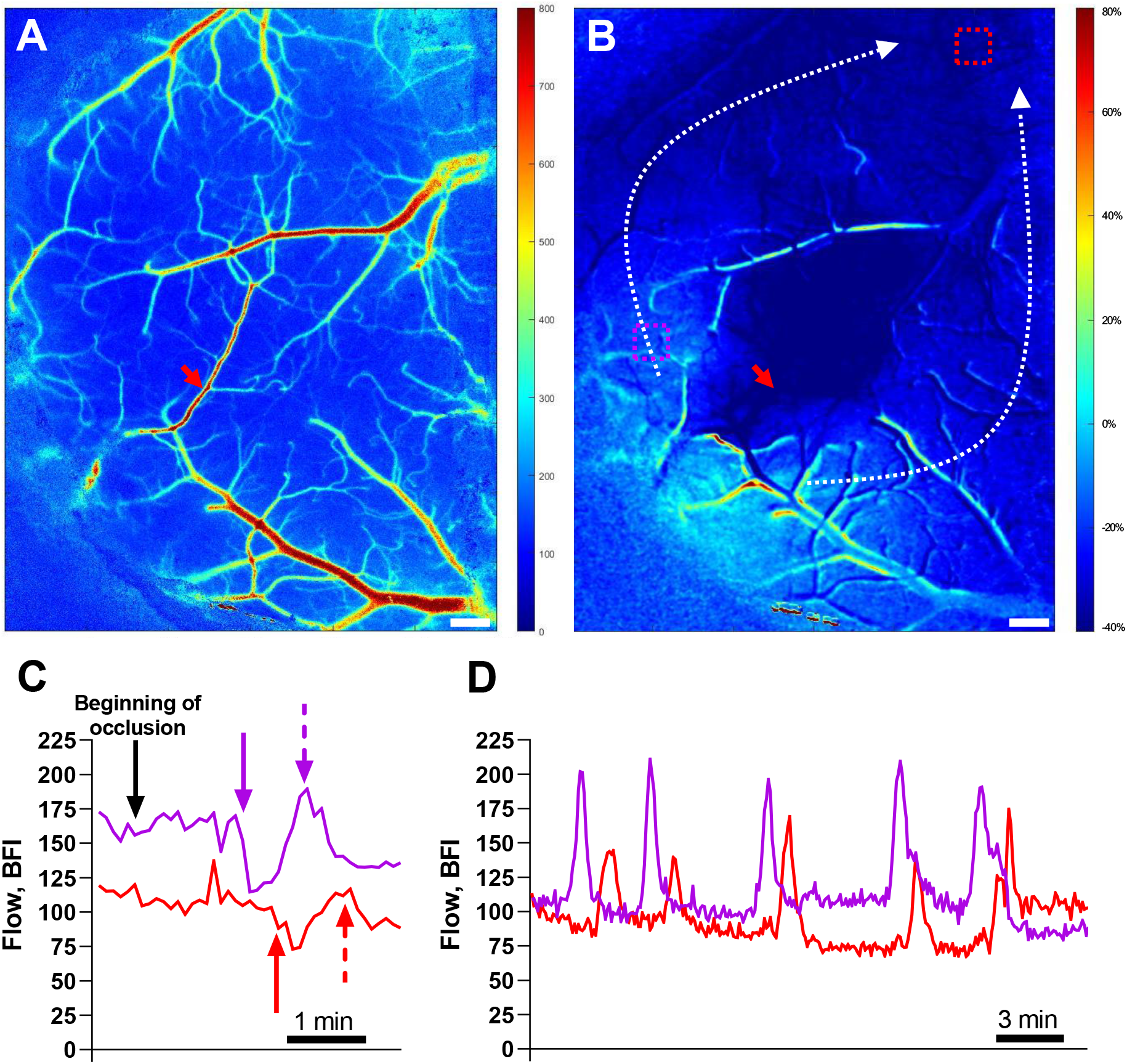
Spreading depressions were elicited by cerebral ischemia. Representative laser speckle contrast image (LSCI) at baseline (A). Ratio of LSCI before vs. during the occlusion identified the ischemic region (B). Red arrows in (A, B) indicate the photothrombosis focus point. Dotted white arrows in (B) show the propagation of spreading depressions that initiated from the lateral part of the penumbral cortex (see also Suppl. Video 1). Two regions of interest (ROIs) were placed just outside the ischemic core (pink dotted box) and in the peri-ischemic area (red dotted box) (B). Bars in (A, B) correspond to 300 μm. Representative traces from these two ROIs at the beginning of occlusion (C) and 25-50 min after initiation of the occlusion (D). The black arrow in (C) indicate the time where the middle cerebral artery occlusion started. Approximately 2 min after the occlusion, the first spreading depression propagated. The solid pink and red arrows in (C) show the beginning of the hypoperfusion whereas the dotted arrows indicate the subsequent hyperperfusion followed by sustained hypoperfusion. During this global hypoperfusion, several spreading depressions propagated and were associated with waves of hyperperfusion (D). The delayed changes in blood flow in the area corresponding to the red ROI is caused by spreading depression propagation. See also Suppl. Table 1 for details about the spreading depressions.

### Reduced neurovascular coupling in peri-ischemic area after ischemic stroke

Neurovascular coupling was assessed at baseline prior to arterial occlusion and 3 and 24h after spontaneous reperfusion. At baseline, 5-sec air-puff whisker stimulation was associated with a localised increase in blood flow in the contralateral whisker sensory cortex (Fig. 3A-B). This neurovascular response was reduced 3 and 24h after reperfusion (Fig. 3B).

**Figure 3.**
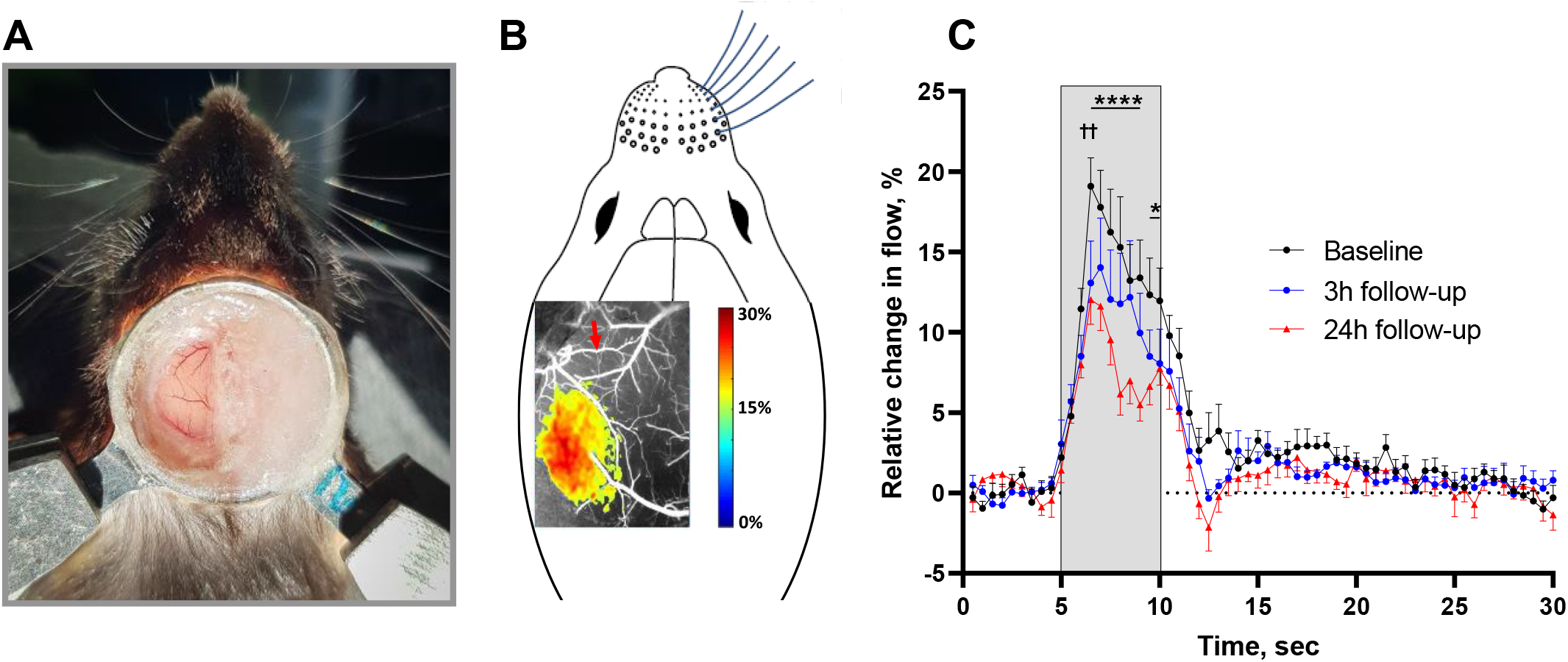
Neurovascular coupling responses in the whisker sensory cortex corresponding to the peri-ischemic area were reduced after cerebral ischemia. Image of the chronic cranial window preparation and the head fixation of the awake mouse 4 wks after surgery (A). Representative laser speckle contrast image illustrating local increase in blood flow in the whisker sensory cortex in response to air-puff whisker stimulation at baseline (B). Red arrow in (B) indicates the artery that later was occluded. The neurovascular coupling response was reduced 3 and 24h after ischemia-reperfusion compared to baseline (C). Grey area in (C) indicate the period when the 5 sec whisker stimulation was performed. Neurovascular responses were compared using two-way ANOVA followed by Bonferroni’s correction for multiple comparison. †† indicates *P* < 0.01 when comparing the neurovascular response at 3h follow-up with baseline. *, **** indicate *P* < 0.05, 0.0001 when comparing the response at 24h follow-up with the response at baseline; *n* = 6.

### Resting perfusion of the peri-ischemic cortex was reduced 3 hours after reperfusion but returned to the baseline value after 24 hours

Results from LSCI (Fig. 4A) showed that perfusion of the cortex downstream from the occluded MCA branch remained suppressed 3 and 24h after reperfusion (Fig. 4B). In the peri-ischemic area, cortical perfusion was also reduced 3h after reperfusion, although to a lesser extent than in the ischemic core (Fig. 4B). At the 24h follow-up, perfusion of the peri-ischemic cortex was similar to that observed at baseline.

**Figure 4.**
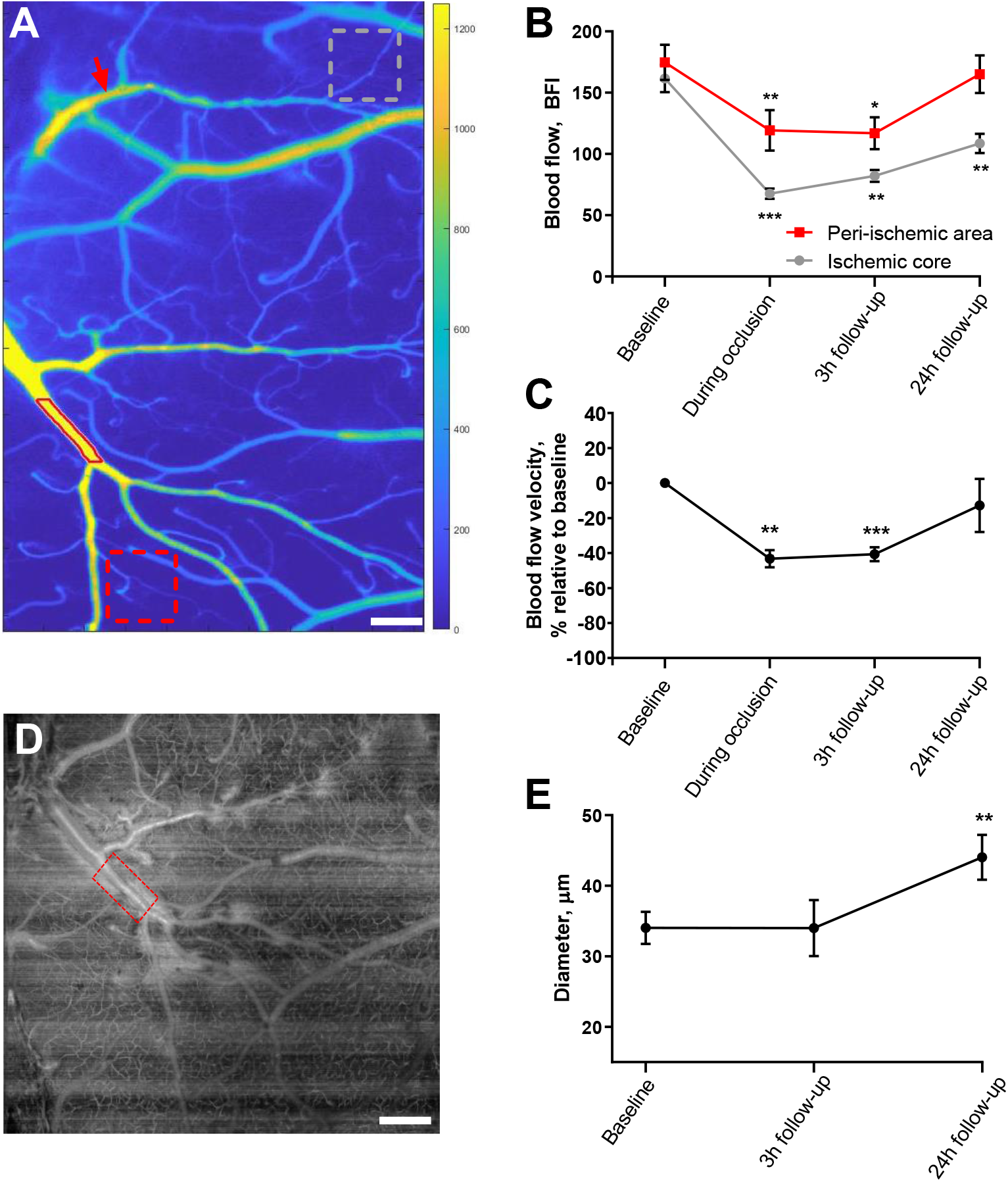
Cerebral perfusion was globally reduced 3 hours after ischemia-reperfusion but returned to baseline after 24 hours. Tissue perfusion was measured by laser speckle contrast imaging (A). Regions of interest for tissue perfusion (grey and back dotted boxes in the ischemic core and peri-ischemic area, respectively) and the vessel segmentation (lumen outlined by red line) of the middle cerebral artery (MCA) branch neighbouring to the occluded artery are shown (A). Red arrow in (A) indicates the focus point for photothrombosis. Tissue perfusion was globally reduced 3h after ischemia-reperfusion in both the ischemic core and the peri-ischemic area (B). At the 24h follow-up, the tissue perfusion of the peri-ischemic area returned to a level similar to the perfusion observed at baseline (B). Blood flow velocity in the MCA branch supplying the peri-ischemic area was reduced during the occlusion of the anterior MCA branch and at the 3h follow-up but was not statistically different from baseline at the 24h follow-up (C). Representative optical coherence tomography (OCT) angiogram (D) showing the segment (red box) of the MCA branch neighbouring to the occluded artery; same MCA segment as shown in (A). Diameter of the neighbouring MCA branch was assessed from this angiogram and revealed vasodilation at the 24h follow-up (E). Bars correspond to 300 (in A) and 400 μm (in D). Data were compared to baseline using mixed-effects analysis followed by Dunnett’s multiple comparisons test. *, **, *** indicate *P* < 0.05, 0.001, 0.0001; *n* = 6.

Blood flow was also assessed in the neighbouring MCA branch of the occluded anterior MCA branch (Fig 4A). Similar to the observed changes in tissue perfusion of the peri-ischemic area, blood flow velocity in this neighbouring artery was reduced during the occlusion and 3h after reperfusion but was not statistically different from baseline at the 24h follow-up (Fig. 4C). Diameter of the neighbouring MCA branch remained unchanged from baseline at the 3h follow-up, but the artery showed vasodilation at the 24h follow-up (Fig. 4D-E).

### Increased flow stalling of the capillary circulation after reperfusion in the peri-ischemic area

OCT angiography (Fig. 5A-F) showed that the number of perfused capillaries was reduced compared to baseline when assessed 3 and 24h after reperfusion (Fig. 5G). Furthermore, the incidence of flow stalling (Suppl. Video 2 and Fig. 5H) and stalling duration (Fig. 5I) were increased 3 and 24h after reperfusion compared to baseline. Consequently, the point prevalence of dynamic capillary flow stalling was increased, i.e., the risk of any capillary being blocked at any given time was increased 3 and 24h after reperfusion compared to baseline (Fig. 5J). The mean duration of each capillary stalling was increased 3h after reperfusion; the mean stalling duration was not statistically different at baseline and at the 24h follow-up (Fig. 5K). The frequency distribution of the mean duration of each capillary stalling is shown in Suppl. Fig. 1.

**Figure 5.**
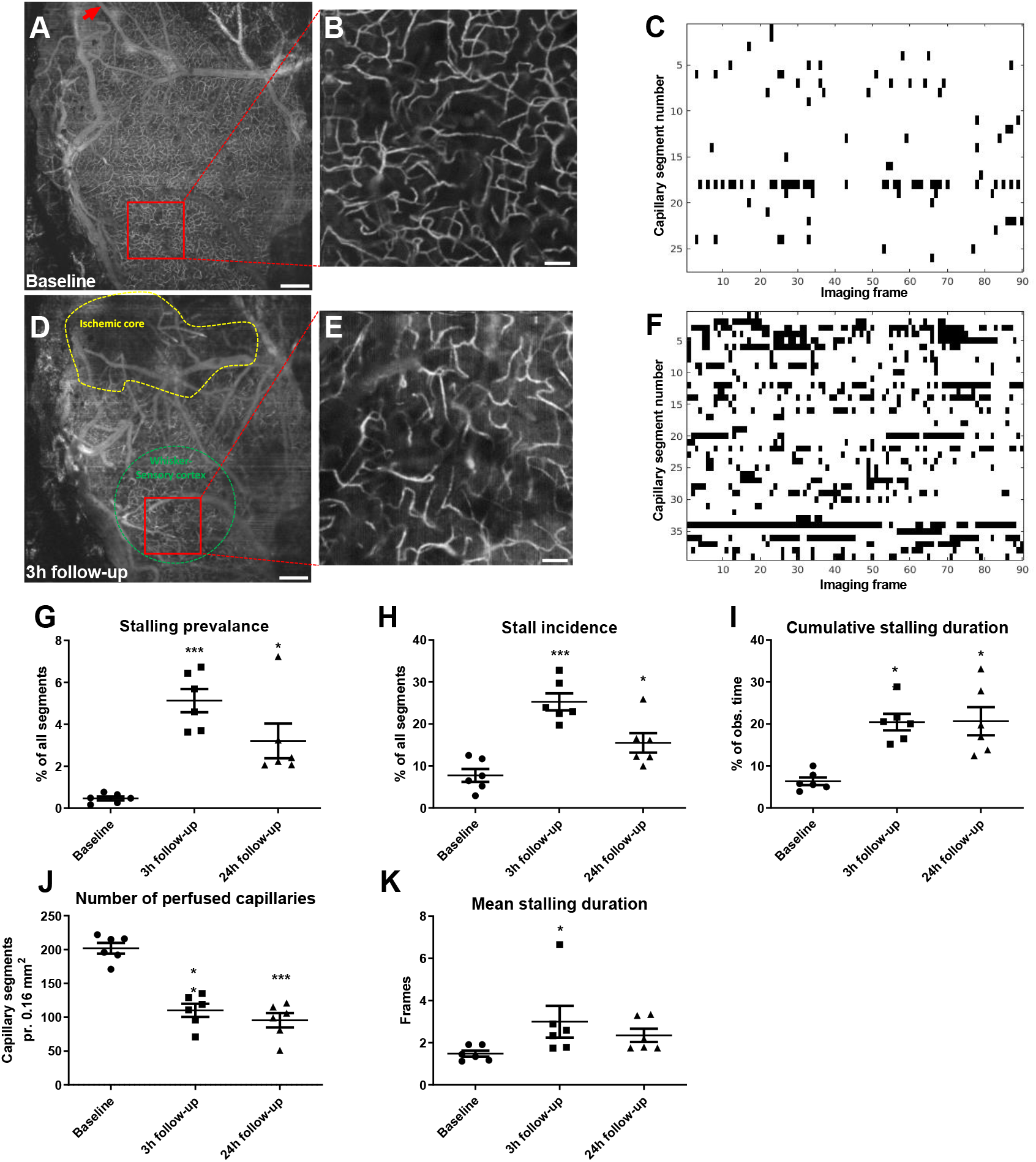
Reduced number of perfused capillaries and increased dynamic capillary flow stalling after ischemia-reperfusion. Representative optical coherence tomography (OCT) images of the affected hemisphere at baseline (A, B) and 3h after reperfusion (D, E). Bars in (A, D) correspond to 300 μm and bars in (B, E) 40 μm. The red arrow in (A) indicates the location of photothrombosis. Yellow and green dotted areas in (D) indicate the ischemic core and the whisker sensory cortex, respectively. The high magnification OCT angiograms (B, E) were used to assess capillary flow stalling (see Suppl. Video 2). Representative stallograms from the same mouse at baseline (C) and 3h after reperfusion (F) showing the timeline of stalling capillary segments through the 90 consecutive images. The number of perfused capillaries was reduced at the 3h follow-up and 24h follow-up compared to baseline (G). The incidence of dynamic flow stalling (H), stalling duration (I), and thus stalling prevalence (J) were increased at the 3h follow-up and 24h follow-up compared to baseline. The mean duration of each stalling event was increased at the 3h follow-up but not statistically different at the 24h follow-up compared to baseline (K). Data were compared to baseline using repeated measures one-way ANOVA followed by Dunnett’s multiple comparisons test. *, **, *** indicate *P* < 0.05, 0.01, 0.001; *n* = 6.

### Capillary blood flow velocity increased 24 hours after reperfusion

Blood flow velocity in the capillary circulation of the peri-ischemic area was assessed by prD-OCT (Fig. 6A). This was done in the same region of interest and depth range as the OCT angiograms used for capillary flow stalling analysis. Capillary flow velocity was increased at the 24h follow-up compared to baseline, whereas there was no difference in flow velocity between 3h follow-up and baseline (Fig. 6B). Mean inner capillary diameter of perfused capillaries was similar at all three time points (4.67 ± 0.04 μm, 4.53 ± 0.12 μm, and 4.69 ± 0.06 μm for baseline, 3 and 24h follow-up, respectively).

**Figure 6.**
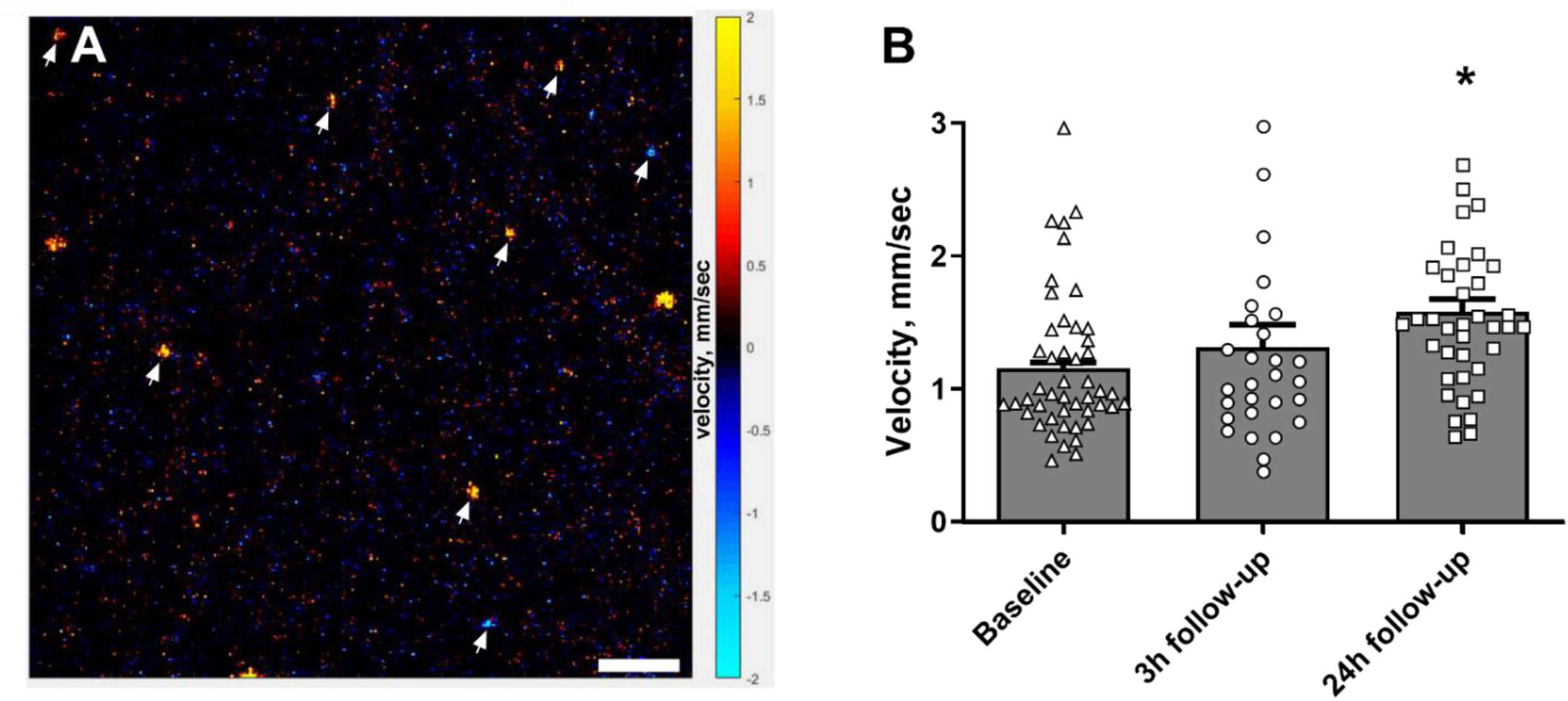
Capillary flow velocity was increased 24 hours after reperfusion. Representative phase-resolved doppler optical coherence tomography image at baseline showing capillaries (white arrows) penetrating the cortex in a vertical direction (A); bar indicates 50 μm, red/yellow and blue/turquoise colours indicate ascending and descending flow, respectively. Average capillary flow velocity was similar 3h after reperfusion but increased at the 24h follow-up compared to baseline (B). Measurements from capillaries for each mouse at each time point were averaged and plotted as a bar graph. Individual capillary velocity measurements from all mice are plotted on top of bar graphs. The weighted average of capillary flow velocity at the two follow-up time points was compared to baseline using repeated measures one-way ANOVA followed by Dunnett’s multiple comparisons test. * indicates *P* < 0.05, *n* = 6.

### Reduced diameter of pericyte-associated capillaries in the peri-ischemic tissue

Capillary diameter was assessed in perfusion-fixed brain tissue (Fig. 7A-B). The diameter of capillaries surrounded by pericyte bodies was narrower in the peri-ischemic tissue compared with corresponding pericyte-surrounded capillaries from the contralateral hemisphere (Fig. 7C). Diameter of capillaries, which were not associated with pericyte bodies, was similar in the peri-ischemic and contralateral hemisphere. Capillary density was not different between the two hemispheres in the perfusion-fixed brains (Fig. 7D).

**Figure 7.**
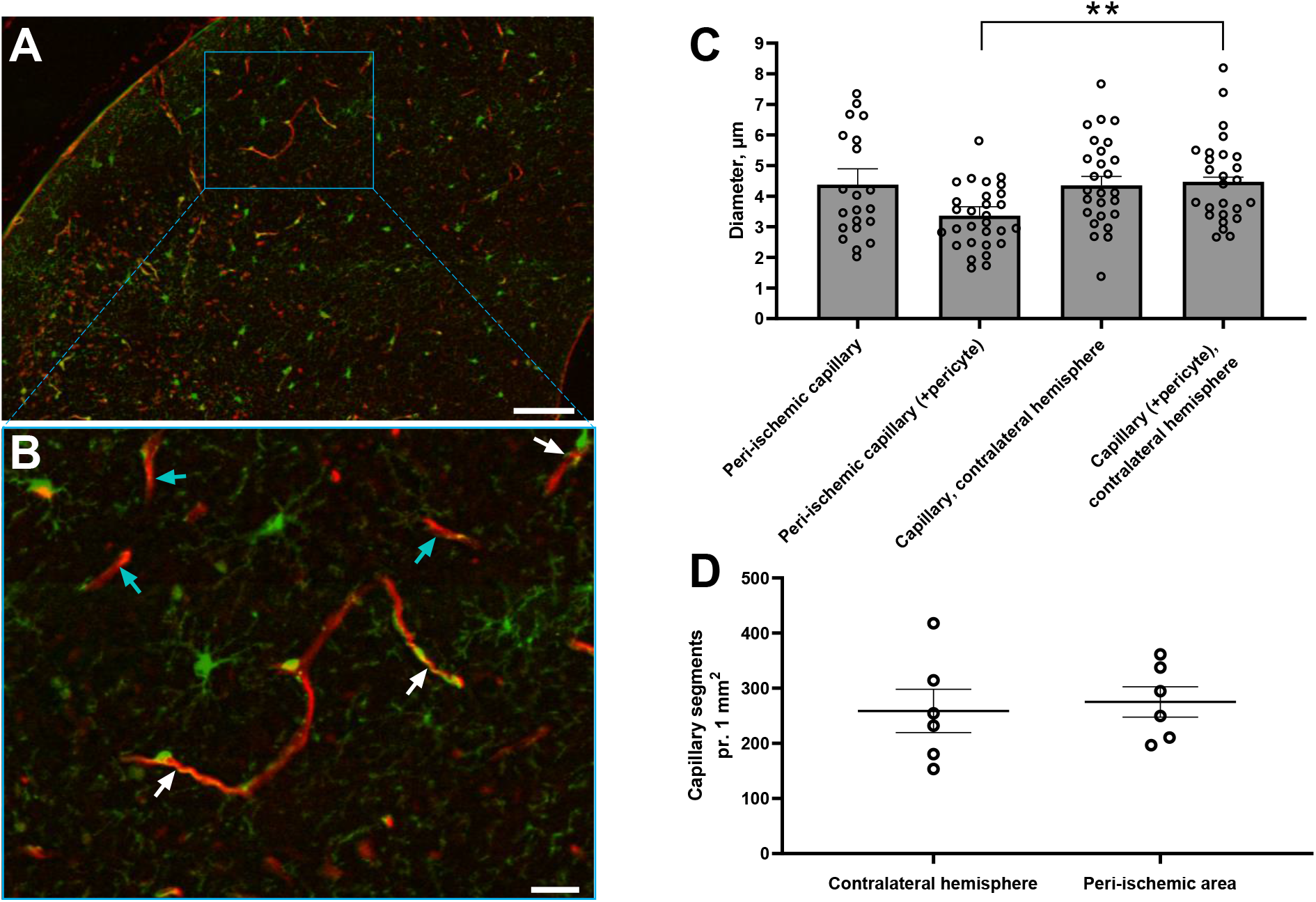
Diameter of capillary segments surrounded by pericyte bodies was reduced in the peri-ischemic cortex. Representative pictures from the peri-ischemic cortex showing PDGFRβ-positive cells in green colour and lectin in red colour in low (A) and high magnification (B). White and turquoise arrows in (B) indicate capillary segments with and without pericyte bodies, respectively. Bar in (A) corresponds to 100 μm and bar in (B) 25 μm. Diameter of capillaries associated with pericyte bodies in the peri-ischemic cortex was reduced compared to the contralateral cortex (C). There was no difference in diameter of capillaries that were not associated with pericyte bodies between the hemispheres. The density of capillary segments was similar in both hemispheres (D). Capillary diameter in peri-ischemic cortex vs. contralateral hemisphere was compared using unpaired t-test. ** indicate *P* < 0.01, *n* = 6.

## Discussion

### Capillary flow heterogeneity and its role in ischemic stroke

In this study, neurovascular coupling in the peri-ischemic area was impaired following reperfusion, associated with constriction of peri-ischemic capillary pericytes. Pericyte constriction was associated with a reduced number of perfused capillaries, although the total number of capillaries in the peri-ischemic area was similar to that observed in the contralateral hemisphere. The reduced capillary capacity was further worsened by increased dynamic flow stalling in the capillaries that were not permanently occluded. Consistent with reduced capillary capacity, overall parenchymal perfusion was reduced in the peri-ischemic area 3h after reperfusion. However, at the 24h follow-up, upstream arterial vasodilation was associated with parenchymal perfusion similar to that observed at baseline, although capillary capacity remained suppressed. Consequently, capillary blood flow velocity was increased.

Unequal blood flow distribution in the capillary bed increases capillary transit time heterogeneity associated with impaired oxygen extraction capability (Jespersen and Østergaard, 2012). Such undesired blood flow distribution, also known as “functional shunting”, is caused by an increasing proportion of erythrocytes passing through the capillary at transit times too short to allow optimal oxygen extraction (Østergaard et al., 2013). Cerebral blood flow distribution in ischemic stroke patients assessed by perfusion-weighted magnetic resonance imaging (MRI) revealed increased capillary transit time heterogeneity in the penumbra (Engedal et al., 2018). Hence, increased capillary transit time heterogeneity may prevent sufficient oxygen delivery to active brain regions mediated through neurovascular coupling in stroke patients. Accordingly, previous BOLD fMRI studies on stroke patients reported impaired oxygen extraction in peri-ischemic areas in response to the performance of simple motor tasks (Krainik et al., 2005; Pineiro et al., 2002). Considering data from the present study, disrupted capillary circulation and thus reduced capillary capacity may underlie disturbances in neurovascular coupling in these patients.

### Disrupted capillary perfusion in peri-ischemic brain regions

Capillary blockage downstream from the occluded artery has been suggested to explain the no-reflow phenomenon in the ischemic core (Korte et al., 2022; Østergaard et al., 2013). There may be several explanations for this outcome, including arterial vasospasm (Guldbrandsen et al., 2021) and blood-brain barrier disruption leading to cerebral edema (Ng et al., 2021). Another possible explanation is the constriction of capillary pericytes with such activity being sustained in ischemia (Hall et al., 2014; Yemisci et al., 2009). In the present study, pericyte constriction associated with microvascular failure in the peri-ischemic area may be initiated by hypoxia. Another possible explanation is that the pericyte constriction was elicited by the multiple spreading depressions that occurred during arterial occlusion. Experimentally induced spreading depression was recently demonstrated to cause constriction of capillary pericytes associated with neurovascular uncoupling in anesthetized mice (Khennouf et al., 2018). In the present study, the first out of multiple spreading depressions was observed within minutes after the targeted MCA branch was occluded. This caused a significant drop in blood flow in the peri-ischemic area. At the 3h follow-up, where a substantially reduced number of perfused capillaries was observed, the perfusion in the peri-ischemic area remained suppressed to a similar level to that observed during the occlusion. This implies that spreading depressions during arterial occlusion led to pericyte constriction and thus microvascular failure in the peri-ischemic area.

Spreading depressions that propagate from the peri-ischemic area during cerebral ischemia were originally described by Branston (Branston et al., 1977). Pre-clinical studies have shown that the shape, distribution, and number of spreading depressions correlate with the expansion of the ischemic core (Binder et al., 2022; Mies et al., 1993; Unekawa et al., 2022). The excessive metabolic workload of the neuronal tissue was speculated to be caused by spreading depression, which might underlie the transition of salvageable penumbral tissue into irreversible ischemia (Mies et al., 1993). Consistent with the present data, another possible explanation is that spreading depression initiates pericyte constriction and thus failure of the capillary circulation in the peri-ischemic area resulting in compromised oxygen extraction and thus ischemia.

### Increased incidence of dynamic capillary flow stalling after reperfusion

In addition to microcirculatory failure, we also observed increased dynamic flow stalling in the peri-ischemic area. Such dynamic stallings were recently reported in penumbra in a pre-clinical model for transient cerebral ischemia (Erdener Ş et al., 2021). Although done in anesthetised mice, the incidence and point prevalence of the stallings were similar to those observed in the present study. Intriguingly, specific pharmacological targeting of Ly6G, a surface protein expressed exclusively in neutrophils in mice, reduced the stalling events (Erdener Ş et al., 2021). A similar reduction in dynamic capillary stalling after anti-Ly6G antibody treatment has been observed in a pre-clinical model for Alzheimer’s disease (Cruz Hernández et al., 2019). This outcome suggests that dynamic flow stalling, at least in part, is caused by adhesion of neutrophils to the capillary wall. In both studies, anti-Ly6G treatment improved the neurological outcome (Cruz Hernández et al., 2019; Erdener Ş et al., 2021). Hence, dynamic capillary flow stalling, which was also observed in the present study, may be a novel target in the treatment of cerebrovascular disease including ischemic stroke.

Some limitations need to be considered in this study. First, although an optimized protocol for photothrombosis was used that limits the occlusion to a specific narrow location, it is possible that a few capillaries right below the targeted artery were also occluded. However, surrounding capillaries and larger vessels are not thrombosed when using this protocol (Sunil et al., 2020). Second, using a mouse model to study cerebrovascular pathology in ischemic stroke has inherent limitations in representing the human state. Thus, it remains to be investigated whether microcirculatory failure in the peri-ischemic area also takes place after ischemic stroke in patients; although this has been suggested by indirect measurements (Engedal et al., 2018). Third, since neuronal activity was not measured, the possibility that neuronal dysfunction after spreading depression may also contribute to impaired neurovascular coupling cannot be excluded. Finally, only the hemisphere where the arterial occlusion took place was examined in vivo. Whether disruption of the microcirculation and thus impaired neurovascular coupling is also affecting the contralateral hemisphere remains to be investigated.

Findings from the present study contribute to an improved understanding of the cerebrovascular changes outside the ischemic core in stroke. Key insights into the vascular events during and after cerebral ischemia are provided, and a novel link between these microcirculatory changes and dysfunctional neurovascular coupling after stroke is proposed. These findings call for future studies on how to enhance overall passage of blood cells through the capillary circulation and thus restore coupling between the neuronal tissue and the vasculature in ischemic stroke patients.

## Acknowledgements

We would like to thank Dr. Dmitry Postnov and Prof. Andrew Dunn for software to acquire and analyse laser speckle data. We also thank John Jiang, Jane Holbæk Rønn, and Anna Bay Nielsen for excellent technical assistance. Finally, we thank Prof. Leif Østergaard and Prof. Christian Aalkjaer for fruitful scientific discussions.

---

**Supplementary Table 1.** Details about spreading depression during arterial occlusion. The first spreading depression in each mouse led to initial hypoperfusion followed by a wave of hyperperfusion and then sustained hypoperfusion. The subsequent spreading depressions during the sustained hypoperfusion were associated with a wave of hyperperfusion. Peak amplitude was calculated as the maximally change in blood flow during spreading depression compared to an average of flow 1 min preceding the spreading depression. See also Fig. 2 for representative traces and Suppl. Video 1. *n* = 6.

| | Mean ± SEM | Lower<br>95% CI | Upper<br>95% CI |
| --- | --- | --- | --- |
| Spreading depression propagation velocity (mm/min) | 4.16 ± 0.21 | 3.61 | 4.71 |
| Time from beginning of occlusion to first spreading depression (sec) | 143.30 ± 32.01 | 61.06 | 225.6 |
| Initial spreading depression hypoperfusion duration (sec) | 36.25 ± 1.25 | 32.27 | 40.23 |
| Amplitude of initial spreading depression hypoperfusion (%) | -33.60 ± 6.01 | -52.71 | -14.48 |
| Spreading depression hyperperfusion duration (sec) | 96.38 ± 6.67 | 79.24 | 113.50 |
| Amplitude of spreading depression hyperperfusion (%) | 72.49 ± 10.59 | 45.28 | 99.70 |
| Number of spreading depressions during 60 min occlusion | 11.33 ± 1.86 | 6.56 | 16.10 |

**Supplementary Figure 1.**
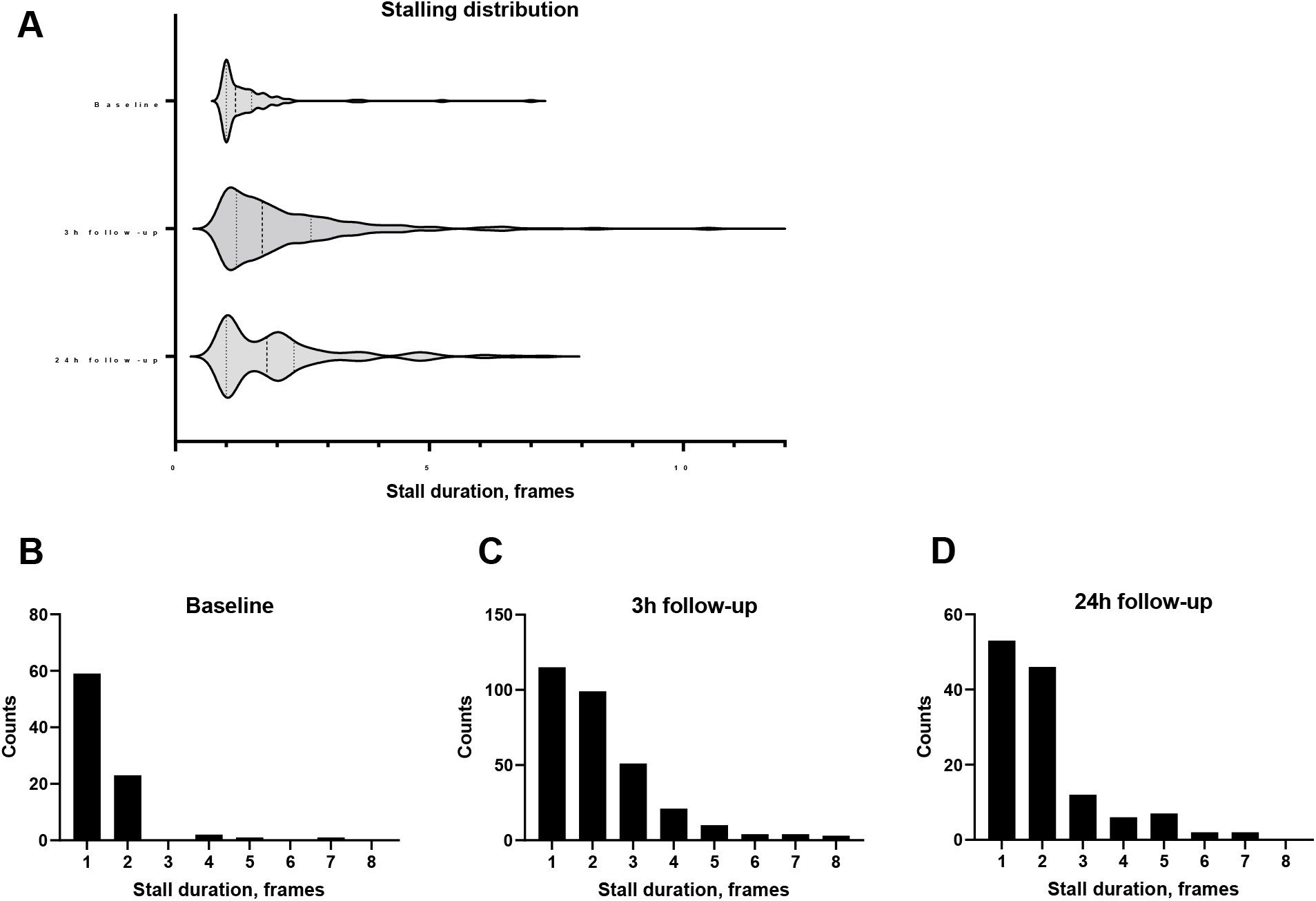
Frequency distribution of mean duration of each capillary stalling. Distribution of stalling durations for all capillary stallings as a violin plot (A). Histograms show the frequency distribution stalling durations at baseline (B), 3h follow-up (C) and 24h follow-up (D). OCT framerate = 0.167 Hz. *n* = 6.

## References

Bandopadhyay, R., et al., 2001. Contractile proteins in pericytes at the blood-brain and blood-retinal barriers. J Neurocytol. 30, 35–44.

Binder, N.F., et al., 2022. Vascular Response to Spreading Depolarization Predicts Stroke Outcome. Stroke. 53, 1386–1395.

Boas, D., Dunn, A., 2010. Laser speckle contrast imaging in biomedical optics. Journal of Biomedical Optics. 15, 011109.

Branston, N.M., Strong, A.J., Symon, L., 1977. Extracellular potassium activity, evoked potential and tissue blood flow. Relationships during progressive ischaemia in baboon cerebral cortex. J Neurol Sci. 32, 305–21.

Cruz Hernández, J.C., et al., 2019. Neutrophil adhesion in brain capillaries reduces cortical blood flow and impairs memory function in Alzheimer’s disease mouse models. Nat Neurosci. 22, 413–420.

D’Esposito, M., Deouell, L.Y., Gazzaley, A., 2003. Alterations in the BOLD fMRI signal with ageing and disease: a challenge for neuroimaging. Nat Rev Neurosci. 4, 863–72.

Engedal, T.S., et al., 2018. Transit time homogenization in ischemic stroke - A novel biomarker of penumbral microvascular failure? J Cereb Blood Flow Metab. 38, 2006–2020.

Erdener Ş, E., et al., 2019. Spatio-temporal dynamics of cerebral capillary segments with stalling red blood cells. J Cereb Blood Flow Metab. 39, 886–900.

Erdener Ş, E., et al., 2021. Dynamic capillary stalls in reperfused ischemic penumbra contribute to injury: A hyperacute role for neutrophils in persistent traffic jams. J Cereb Blood Flow Metab. 41, 236–252.

Erdener, Ş.E., Küreli, G., Dalkara, T., 2022. Contractile apparatus in CNS capillary pericytes. Neurophotonics. 9, 021904.

Fernández-Klett, F., et al., 2010. Pericytes in capillaries are contractile in vivo, but arterioles mediate functional hyperemia in the mouse brain. Proc Natl Acad Sci U S A. 107, 22290–5.

Guldbrandsen, H.O., et al., 2021. Does Src Kinase Mediated Vasoconstriction Impair Penumbral Reperfusion? Stroke. 52, e250–e258.

Hall, C.N., et al., 2014. Capillary pericytes regulate cerebral blood flow in health and disease. Nature. 508, 55–60.

Hill, R.A., et al., 2015. Regional Blood Flow in the Normal and Ischemic Brain Is Controlled by Arteriolar Smooth Muscle Cell Contractility and Not by Capillary Pericytes. Neuron. 87, 95–110.

Jespersen, S.N., Østergaard, L., 2012. The roles of cerebral blood flow, capillary transit time heterogeneity, and oxygen tension in brain oxygenation and metabolism. J Cereb Blood Flow Metab. 32, 264–77.

Khennouf, L., et al., 2018. Active role of capillary pericytes during stimulation-induced activity and spreading depolarization. Brain : a journal of neurology. 141, 2032–2046.

Kisler, K., et al., 2017. Pericyte degeneration leads to neurovascular uncoupling and limits oxygen supply to brain. Nat Neurosci. 20, 406–416.

sKorte, N., et al., 2022. The Ca2+-gated Cl-channel TMEM16A amplifies capillary pericyte contraction reducing cerebral blood flow after ischemia. bioRxiv. 2022.02.03.479031.

Krainik, A., et al., 2005. Regional impairment of cerebrovascular reactivity and BOLD signal in adults after stroke. Stroke. 36, 1146–52.

Lin, W.H., et al., 2011. Impaired neurovascular coupling in ischaemic stroke patients with large or small vessel disease. Eur J Neurol. 18, 731–6.

Longden, T.A., et al., 2017. Capillary K+-sensing initiates retrograde hyperpolarization to increase local cerebral blood flow. Nature Neuroscience. 20, 717–726.

McDowell, K.P., et al., 2021. VasoMetrics: unbiased spatiotemporal analysis of microvascular diameter in multi-photon imaging applications. Quant Imaging Med Surg. 11, 969–982.

Mies, G., Iijima, T., Hossmann, K.A., 1993. Correlation between peri-infarct DC shifts and ischaemic neuronal damage in rat. Neuroreport. 4, 709–11.

Ng, F.C., et al., 2021. Microvascular Dysfunction in Blood-Brain Barrier Disruption and Hypoperfusion Within the Infarct Posttreatment Are Associated With Cerebral Edema. Stroke. Strokeaha121036104.

Nippert, A.R., Biesecker, K.R., Newman, E.A., 2018. Mechanisms Mediating Functional Hyperemia in the Brain. Neuroscientist. 24, 73–83.

Østergaard, L., et al., 2013. The role of the cerebral capillaries in acute ischemic stroke: the extended penumbra model. J Cereb Blood Flow Metab. 33, 635–48.

Pineiro, R., et al., 2002. Altered hemodynamic responses in patients after subcortical stroke measured by functional MRI. Stroke. 33, 103–9.

Postnov, D.D., Tuchin, V.V., Sosnovtseva, O., 2016. Estimation of vessel diameter and blood flow dynamics from laser speckle images. Biomed Opt Express. 7, 2759–68.

Sigler, A., Goroshkov, A., Murphy, T.H., 2008. Hardware and methodology for targeting single brain arterioles for photothrombotic stroke on an upright microscope. J Neurosci Methods. 170, 35–44.

Srinivasan, V.J., et al., 2010. Rapid volumetric angiography of cortical microvasculature with optical coherence tomography. Opt Lett. 35, 43–5.

Staehr, C., et al., 2019. Smooth muscle Ca(2+) sensitization causes hypercontractility of middle cerebral arteries in mice bearing the familial hemiplegic migraine type 2 associated mutation. J Cereb Blood Flow Metab. 39, 1570–1587.

Staehr, C., et al., 2020. Abnormal neurovascular coupling as a cause of excess cerebral vasodilation in familial migraine. Cardiovasc Res. 116, 2009–2020.

Sunil, S., et al., 2020. Awake chronic mouse model of targeted pial vessel occlusion via photothrombosis. Neurophotonics. 7, 015005.

Tang, J., et al., 2017. Capillary red blood cell velocimetry by phase-resolved optical coherence tomography. Opt Lett. 42, 3976–3979.

Tang, J., et al., 2019. Normalized field autocorrelation function-based optical coherence tomography three-dimensional angiography. J Biomed Opt. 24, 1–8.

Unekawa, M., et al., 2022. Close association between spreading depolarization and development of infarction under experimental ischemia in anesthetized male mice. Brain Research. 1792, 148023.

Yemisci, M., et al., 2009. Pericyte contraction induced by oxidative-nitrative stress impairs capillary reflow despite successful opening of an occluded cerebral artery. Nat Med. 15, 1031–7.

